# RUBAT Studio: A Unified Workbench for Multichannel Bioacoustic Data Acquisition

**DOI:** 10.64898/2026.03.01.708807

**Authors:** Ravi Umadi

## Abstract

1. High-resolution ultrasonic recording is central to modern bioacoustics, behavioural ecology, and passive acoustic monitoring. Yet, implementing flexible, multichannel acquisition systems with real-time playback, monitoring and experimental control remains technically demanding. Existing solutions often require bespoke code development or expensive proprietary systems, limiting experimental accessibility and reproducibility.
2. I present RUBAT Studio (v4.0), an integrated software platform for multichannel ultrasonic recording, real-time heterodyne monitoring for experimental control and data acquisition in laboratory and field environments. The system supports high sample rates, configurable channel routing, automated triggering, retrospective ring-buffer capture, calibration-aware visualisation, and modular signal-processing integration. Its architecture is designed for extensibility, enabling synchronised acquisition across multiple microphones and compatibility with custom hardware configurations.
3. Field and laboratory testing with devices ranging from USB microphones to analogue microphones via professional audio interfaces confirmed stable multichannel acquisition at sampling rates up to 384 kHz over multi-hour sessions without crashes or artefacts. Real-time heterodyne monitoring, dual-channel live spectrograms, and calibrated sound-pressure-level display provided immediate acoustic and visual feedback throughout data collection. The platform enables complex experimental paradigms, including spatial localisation studies, closed-loop playback experiments, active-sensing investigations, and array-based behavioural assays, without requiring specialised knowledge of audio systems.
4. By lowering the technical barrier to high-performance audio recording and experimental control, RUBAT Studio expands the methodological toolkit available to behavioural ecologists and bioacousticians. The platform facilitates rigorous, scalable and reproducible acoustic research designs, enabling experiments that were previously technically prohibitive and thereby advancing the study of animal communication, spatial hearing and active sensing.

## 1 INTRODUCTION

Passive acoustic monitoring has become an indispensable tool for biodiversity assessment and conservation biology (Laiolo, 2010; Sugai et al., 2019). By capitalising on the sounds animals produce for orientation, communication, and foraging, bioacousticians can survey populations, describe soundscapes, and track ecological change non-invasively at scales impractical with traditional observation (Pijanowski et al., 2011; Blumstein et al., 2011). The approach is especially powerful for taxa with stereotyped, species-specific vocalisations, enabling acoustic identification and automated classification (Stowell, 2022; Aodha et al., 2022). Across study systems from birdsong and insect stridulation to cetacean communication and bat echolocation, multichannel recordings from spatially distributed microphone arrays have become increasingly important. Array-based approaches enable sound-source localisation, spatial separation of simultaneously vocalising individuals, and estimation of absolute source levels—capabilities fundamental to behavioural experiments and ecological surveys, yet unattainable with single-channel recordings (Blumstein et al., 2011; Verreycken et al., 2021).

Bats (Chiroptera) exemplify both the promise and the technical demands of multichannel bioacoustic recording. Their echolocation calls span roughly 10 to up to 200 kHz, are emitted at high repetition rates, and carry species-diagnostic spectro-temporal signatures (Jones & Holderied, 2007; Fenton et al., 2016). Acoustic surveys have consequently become a primary method for bat diversity inventories, population monitoring, and behavioural studies (Zamora-Gutierrez et al., 2021; Obrist et al., 2010). More broadly, conservation and ecological monitoring increasingly demand source separation and broad-taxonomic classification from multichannel recordings across the full audible and ultrasonic spectrum (Blumstein et al., 2011; Lin & Tsao, 2020; Izadi et al., 2020; Rhinehart et al., 2020; Tolkova & Klinck, 2022).

These requirements highlight a pronounced gap between available software tools and the recording capabilities needed for multichannel bioacoustic research. Hardware options have expanded considerably: affordable autonomous recorders such as AudioMoth (Hill et al., 2019), AURITA (Beason et al., 2019), Solo (Whytock & Christie, 2017), and open-source multichannel platforms such as BATSY4-PRO (Umadi, 2025b) and ultrasound detector-recorders for surveys such as ESPERDYNE (Umadi, 2026a) serve a range of field scenarios. On the analysis side, deep-learning classifiers (Aodha et al., 2022), inspection tools such as BatReviewer (Umadi, 2025a), and general-purpose software (e.g. Raven Pro, Sonic Visualiser) facilitate downstream processing. However, the recording stage itself—the critical bridge between hardware and analysis—remains dominated by proprietary vendor-locked solutions or general-purpose audio workstations not designed for bioacoustic data collection.

Commercial software such as Avisoft RECORDER provides comprehensive real-time features— live spectrograms, heterodyne monitoring, threshold-triggered acquisition—but is optimised for proprietary hardware, Windows-only, and expensive. Kaleidoscope (Wildlife Acoustics) is similarly confined to vendor-specific recorders. PAMGuard (Gillespie et al., 2009) is a notable open-source platform for cetacean monitoring that supports multichannel localisation but requires considerable configuration expertise. General-purpose digital audio workstations (e.g. REAPER, Audacity) offer flexible recording but lack real-time heterodyne monitoring, ultrasonic spectrograms, calibrated sound-pressure-level display, and threshold-triggered acquisition. Other tools, such as AudioMoth Live, offer excellent starting points for live monitoring and recording with external hardware, but are limited by the number of channels and hardware access (a feature-level comparison with existing tools is provided in Table 1).

**Table 1.** Comparison of RUBAT Studio with commonly used acoustic recording and analysis software. Feature support: ✓ = yes, part = partially or indirectly supported (e.g. via plugins or OS-level routing), – = not supported. Platform codes: W = Windows, M = macOS, L = Linux.

| Software | OS | Open/Free | Multi-ch | Het | PT | Live Spec | Pre/Post | Auto | HW-free |
| --- | --- | --- | --- | --- | --- | --- | --- | --- | --- |
| **RUBAT Studio** | WM | ✓ | ✓ | ✓ | ✓ | ✓ | ✓ | ✓ | ✓ |
| Avisoft RECORDER | W | – | ✓ | ✓ | ✓ | ✓ | ✓ | ✓ | part<sup>a</sup> |
| BatSound / Touch | W | – | – | ✓<sup>b</sup> | ✓ | ✓ | ✓ | ✓ | part<sup>c</sup> |
| AudioMoth Live | WML | ✓ | – | ✓ | ✓ | ✓ | ✓ | ✓ | ✓ |
| Kaleidoscope Pro | WM | – | – | – | – | part | – | – | – |
| Raven Pro | WM | – | ✓ | – | part | ✓ | part | – | ✓ |
| PAMGuard | WML | ✓ | ✓ | – | part | ✓ | ✓ | ✓ | ✓ |
| REAPER | WML | – | ✓ | – | ✓ | part<sup>d</sup> | – | – | ✓ |
| Audacity | WML | ✓ | part | – | part | part | – | – | ✓ |

**Columns.** Open = open-source; Multi-ch = multichannel recording ( $\geq 4$ channels); Het = live heterodyne monitoring; PT = real-time passthrough monitoring; Live Spec = real-time spectrogram during acquisition; Pre/Post = pre- and/or post-trigger clip capture; Auto = threshold-triggered automatic recording; HW-free = operates with generic (non-vendor) audio hardware.
<sup>a</sup> Works with third-party ASIO devices but primarily bundled with Avisoft UltraSoundGate hardware.
<sup>b</sup> Virtual heterodyne mode on recorded files; BatSound Touch provides real-time heterodyne. <sup>c</sup> Compatible with any Windows sound card; primarily marketed with Pettersson USB microphones. <sup>d</sup> Via third-party plugins (e.g. JSFX Spectrograph); not a built-in feature.

As a result, researchers assembling multichannel setups with third-party audio interfaces and calibrated microphones must typically resort to custom scripts (Baier et al., 2019; Geberl et al., 2015; Linnenschmidt & Wiegrebe, 2016; Ma et al., 2025), vendor-specific utilities (Corcoran & Conner, 2014; Eitan et al., 2022; de Framond et al., 2023; Jakobsen et al., 2025), or platform-dependent system settings to configure device parameters and manage recording workflows— sometimes requiring separate hardware for monitoring and acquisition (Haron et al., 2026). This fragmentation is compounded by the inherent complexity of multichannel acquisition: device enumeration, sample-rate negotiation, buffer management, and channel routing are handled through OS-specific audio subsystems (Core Audio on macOS, WASAPI/ASIO on Windows) whose configuration interfaces vary across platforms and devices. Many established laboratories already possess high-value equipment—calibrated microphone arrays, custom preamplifiers, multi-port audio interfaces—that must be operated from a host computer, requiring software that exposes and controls all device-level parameters through a single coherent interface.

Here I present RUBAT Studio (v4.0)^1^ (**R**ealtime **U**nified **B**io**A**coustics **T**ool), an open-source, hardware-agnostic multichannel recording environment that addresses these limitations. RUBAT consolidates device management, multichannel recording, real-time visual and auditory monitoring, threshold-triggered acquisition, and signal-level calibration into a unified graphical interface, surfacing all relevant audio-device parameters—normally dispersed across proprietary utilities and OS settings—in a single control panel. The software dynamically discovers sampling rates supported by each connected device (44.1 to 768 kHz), supports up to 64 input channels, and offers three recording paradigms—manual tap with pre/post-trigger, continuous streaming, and automatic threshold-triggered capture (Section 2.5). Real-time spectrograms and waveform displays coupled with selectable heterodyne and passthrough monitoring (Section 2.7) provide continuous acoustic and visual feedback throughout data collection. RUBAT runs on macOS and Windows with any standard USB or built-in audio device and is freely available as compiled standalone executables that require only the free MATLAB Runtime.

## 2 DESIGN AND IMPLEMENTATION

RUBAT Studio (v4.0) (Figure 1) was developed in MATLAB (R2023b and later) using the Audio Toolbox for low-latency device access and streaming. The application is implemented as a single top-level function enclosing approximately 40 nested routines that share a lexical closure over a central state structure and all user interface handles. No class hierarchy or object-oriented decomposition is employed; the entire application resides in a single self-contained source file.

**Figure 1.**
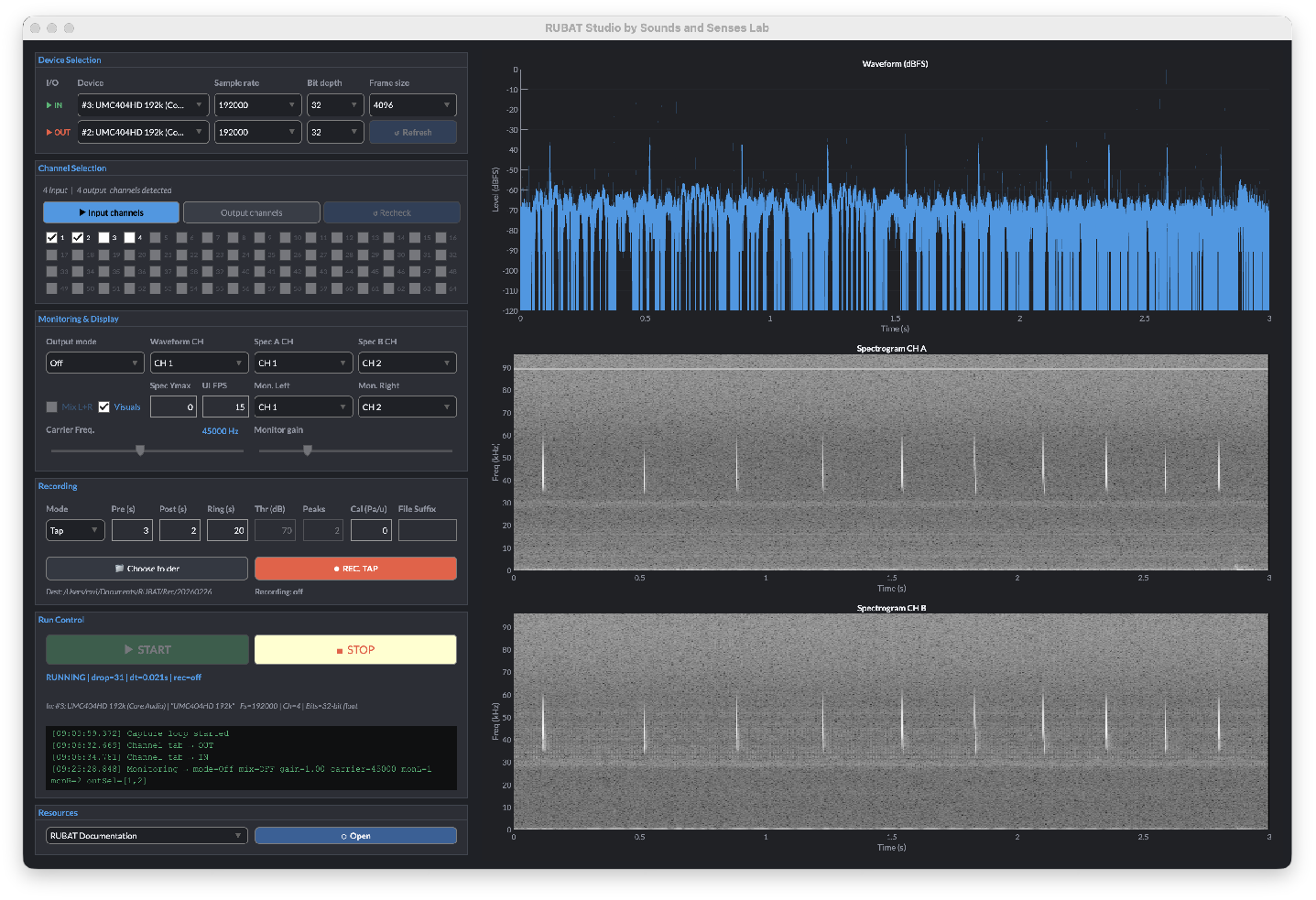
The RUBAT Studio (v4.0) graphical interface. The left panel provides controls for input and output device selection, channel assignment with a soft limit of 64 channels, sampling rate and bit-depth configuration, monitoring mode (off, passthrough, or heterodyne), recording mode (tap, continuous, or auto), and calibration. The right panel displays the real-time waveform (top) and dual live spectrograms (bottom) for two independently selectable channels. An embedded log console at the bottom of the control panel records timestamped operational events for diagnostics and reproducibility.

### 2.1 User interface design

The graphical interface (Figure 1) is rendered using MATLAB’s `uifigure` framework in a darkthemed layout (1 500 × 980 px). The root grid is divided into a fixed-width left panel (520 px) containing six vertically stacked control sections—device selection, channel selection, monitoring and display, recording, run control, and external resources—and a flexible right panel hosting the waveform and dual spectrogram axes. Channel selection is presented through a simulated tab interface supporting up to 64 channels (4 rows × 16 columns), with channels exceeding the detected device maximum greyed out.

All device selection controls are disabled during active streaming to prevent mid-run configuration errors. Keyboard shortcuts (`s/S` for start/stop, `t` for record/arm, `x` for stop continuous) complement the button-driven interface for rapid field operation. Contextual tooltips are provided for all interactive controls.

All operations are logged with timestamps and rendered in an embedded HTML console, colour-coded by severity (green for normal, yellow for warnings, orange for errors). The log auto-scrolls and retains up to 400 entries, supporting transparent diagnostics and reproducibility during deployments.

### 2.2 Software architecture

A flat structure (`S`) holds all runtime state, including device registries, buffer pointers, recording flags, auto-trigger accumulators, heterodyne phase, circular display buffers, and frame-timing counters. User interactions—such as device selection, monitoring mode changes, and recording parameter adjustments—modify this structure through widget callbacks. The real-time acquisition path is driven by a synchronous `while` loop (`captureLoop`) that blocks on the `audioDeviceReader` object at the audio frame rate. Graphical updates are dispatched at a configurable refresh rate (default 15 Hz), thereby decoupling visual rendering from data capture and ensuring that user interface operations cannot interrupt the audio stream. Functions that do not require loop access—including WAV streaming, device enumeration utilities, and UI factory helpers—are implemented as local functions outside the main loop.

A prime consideration for developing this software for elaborate experimental setups is depicted in Figure 2, illustrating a representative multi-room laboratory scenario in which a single host computer running RUBAT serves as the central recording terminal for several concurrent experimental setups. Because RUBAT treats each connected audio interface as an independently selectable device, the operator can switch between rooms—and between recording and monitoring hardware—entirely from software, without physical recabling.

**Figure 2.**
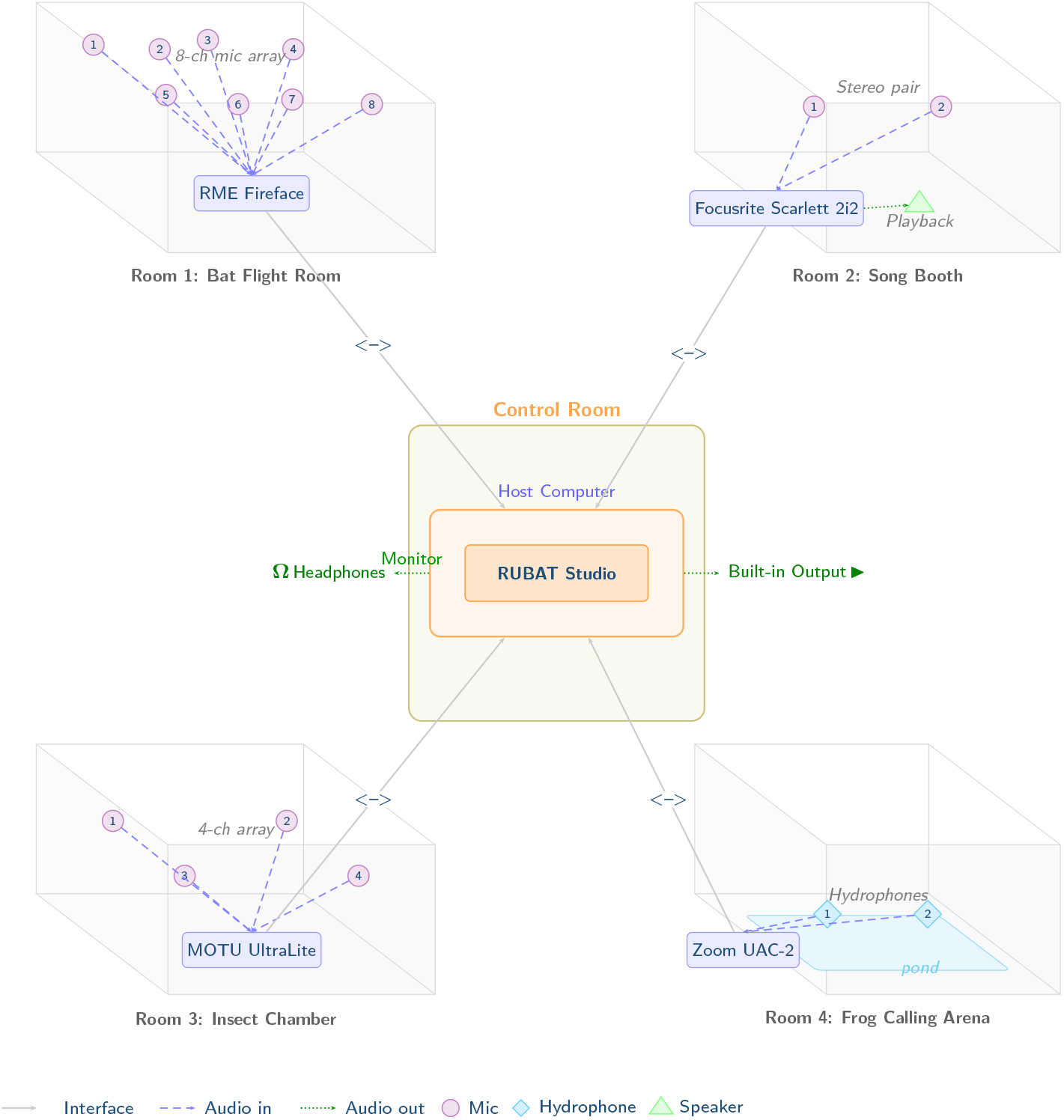
Multi-room laboratory use case. Four experimental rooms—a bat flight room with an eight-channel ultrasonic array (Room 1), a birdsong playback booth with a stereo pair (Room 2), an insect chamber with a four-channel array (Room 3), and a frog calling arena with two hydrophones (Room 4)—surround a central operator control room, each equipped with an independent USB audio interface connected to a single host computer. RUBAT Studio runs as a central recording terminal: the operator selects the desired input device from the interface dropdown, configures channels and sampling rate, and begins recording—without disconnecting or recabling hardware. Cross-device routing allows the input (recording) and output (monitoring) devices to differ; for example, recording from the RME Fireface in Room 1 while monitoring through the computer’s built-in headphone output, or routing playback stimuli through the Focusrite in Room 2. The mentioned commercial audio interfaces are for demonstrative purposes only.

### 2.3 Audio device management and probing

Device enumeration is performed through `audiodevinfo` (PortAudio-backed) in conjunction with the Audio Toolbox classes `audioDeviceReader` and `audioDeviceWriter`. Input and output devices are maintained in separate registries, each entry storing the device name, sorting the lists with properties.

Device probing employs a brute-force combinatorial search over device-name variants, sampling rates, and channel counts. For each candidate combination, a reader (or writer) object is instantiated, invoked once, and the negotiated parameters are read back. The search order prioritises the user-requested rate, followed by a descending sequence of standard rates, and the channel search likewise descends from 64 to 1. Per-device supported sample rates and bit depths are additionally queried from the `audiodevinfo` structure fields and used to populate the corresponding user interface dropdowns dynamically. Every probe attempt is logged with its full parameter set and outcome to facilitate transparent diagnostics.

For same-device duplex configurations, the output sampling rate is locked to the input rate and the writer is opened synchronously, because PortAudio commits a single shared buffer size per device. When monitoring is disabled, silence frames are written each cycle to prevent output buffer starvation. All device handles are explicitly released upon stream termination, with a platform-specific delay on macOS to accommodate Core Audio’s asynchronous port deallocation.

Figure 3 provides an overview of the signal-flow architecture and the three parallel processing paths within the capture loop. The subsections that follow describe each stage in detail.

**Figure 3.**
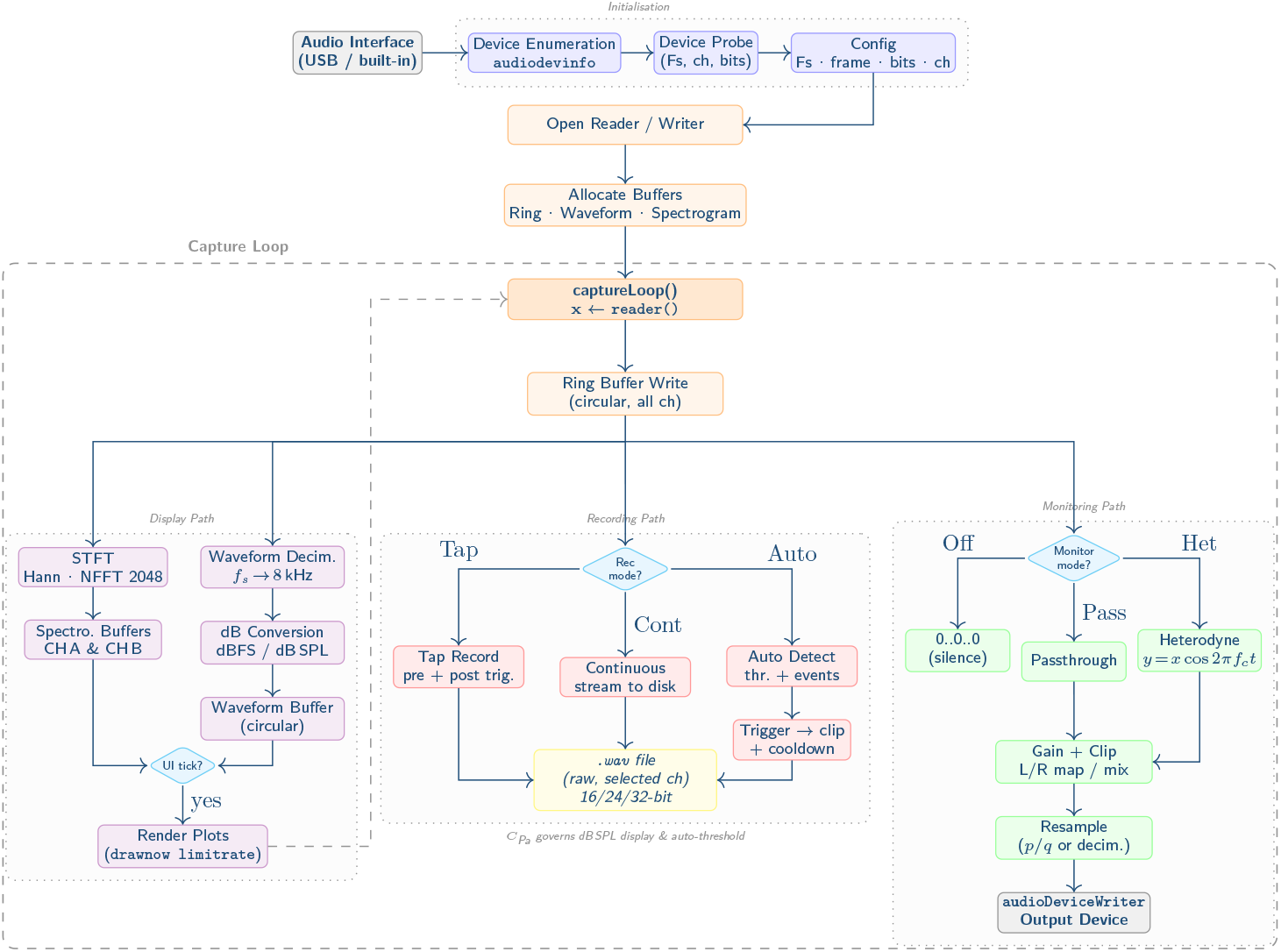
Signal-flow and operational architecture of RUBAT Studio. Rounded rectangles represent processing stages; diamonds are conditional dispatchers. The three parallel paths—display (violet), recording (red), and monitoring (green)—operate independently within the synchronous capture loop. Grey dashed outline delimits the real-time loop; dotted groups indicate functional subsystems. Hardware nodes are shown in grey. The recording path writes only the raw microphone-level signal; monitoring gain, heterodyne processing, and calibration scaling are never applied to saved files.

### 2.4 Sampling framework and buffering

RUBAT Studio discovers the sampling rates supported by each connected audio device at runtime and populates the rate-selection dropdown accordingly. The default set of offered rates spans 44.1, 48, 88.2, 96, 176.4, 192, 352.8, 384, 705.6, and 768 kHz; on platforms where the operating-system audio subsystem reports per-device capabilities (e.g. Windows WASAPI), the list is narrowed to confirmed rates. The default rate is 192 kHz, chosen to accommodate the ultrasonic frequency range relevant to bat echolocation. The frame (buffer) size is user-selectable from 256 to 16 384 samples (default 4 096); at the default settings this corresponds to approximately 21.3 ms per frame.

A circular ring buffer stores a configurable duration of recent samples (default 20 s, with an enforced minimum of the pre-trigger duration plus 0.5 s). The buffer is allocated in single-precision floating point across all available input channels at stream start-up. Incoming frames are written continuously using a classic circular overwrite with wrap-around indexing (Figure 4). Pre-trigger recording is implemented by extracting the required number of samples retrospectively from the ring buffer (see Section 2.5); channel selection is applied at read time rather than at write time, so the full device channel matrix is always preserved.

**Figure 4.**
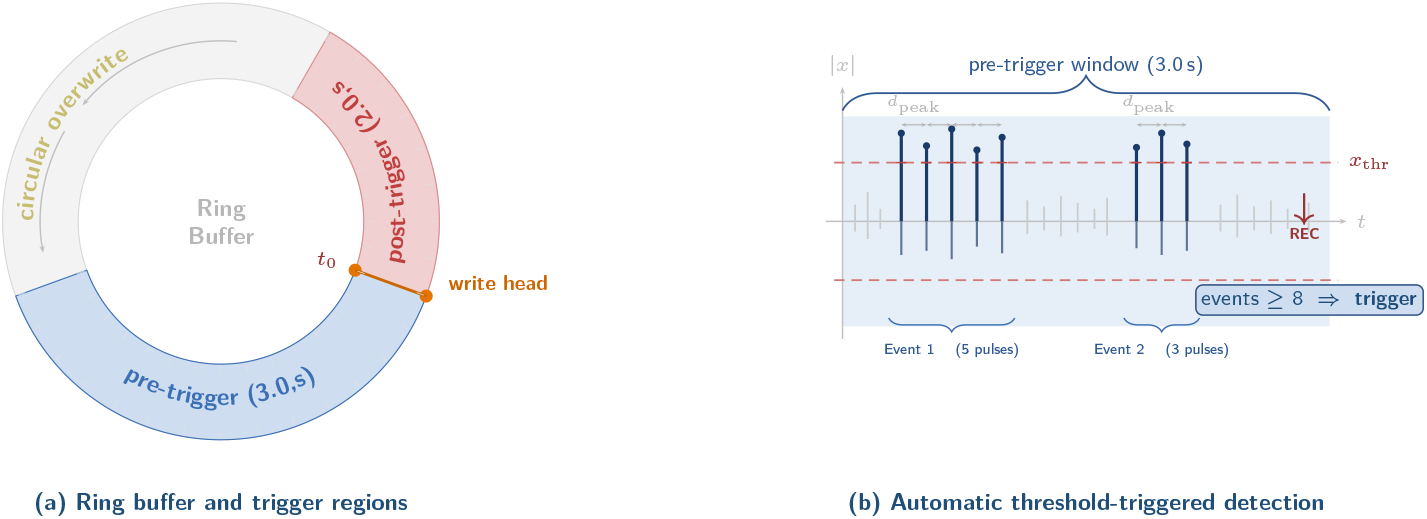
Ring buffer and automatic recording trigger. **(a)** The circular ring buffer continuously overwrites incoming audio frames. When a trigger event occurs at time *t*_0_, the pre-trigger segment (blue) is extracted retrospectively from the buffer and the post-trigger segment (red) is captured from incoming frames, together forming the saved clip. **(b)** Auto-trigger detection: dark vertical lines represent acoustic peaks; those exceeding the amplitude threshold *x*_thr_ (dashed red line) are counted as threshold crossings, while shorter sub-threshold peaks (grey) are ignored. Consecutive crossings whose inter-peak distance *d*_peak_ is less than a configurable minimum (default 3 ms) are merged; crossings separated by more than *d*_peak_ are counted independently. Recording is triggered when the cumulative number of threshold crossings within the pre-trigger window (blue shading, matching the ring-buffer pre-trigger region) meets or exceeds the user-defined minimum—set to 8 in this example, after which the detector fires (red arrow). The default minimum is 2, which is deliberately moderate: even two threshold crossings suffice to distinguish a genuine acoustic event from isolated transient noise. The peak distance *d*_peak_ is fixed at 3 ms and is not adjustable from the graphical interface, but can be modified in the source code to suit different target taxa.

For waveform display, the input signal is decimated from the device sampling rate to a plotting rate of 8 kHz and stored as decibel values (dBFS or dB SPL, depending on calibration state) in a separate circular display buffer. Real-time performance diagnostics—including measured frame intervals and audio drop event counts—are monitored internally to detect timing irregularities during acquisition.

### 2.5 Recording modes

Three recording paradigms are implemented. In all modes, saved audio data consist of the raw microphone-level signal; monitoring gain, heterodyne processing, and calibration scaling are never applied to recorded files.

#### Tap mode

A manual trigger captures a clip consisting of user-defined pre-trigger (default 3.0 s) and post-trigger (default 2.0 s) durations. Upon activation, the pre-trigger segment is extracted from the ring buffer, after which incoming frames are appended until the post-trigger sample count is reached. Only the user-selected input channels are retained in the output file.

#### Continuous mode

Audio is streamed directly to disk via a lightweight custom WAV writer until manually stopped. The ring buffer pre-trigger segment is written at the start of the stream, ensuring that acoustic events immediately preceding activation are captured. Pre-trigger, post-trigger, and ring-buffer duration controls are disabled in this mode. This approach supports extended unattended field deployments without imposing memory constraints.

#### Auto mode

A lightweight threshold-based detector automatically triggers recording. The user-selected waveform channel is evaluated in real time using the following procedure. First, all sample indices with absolute amplitudes exceeding the threshold are identified as threshold crossings. Consecutive crossings whose temporal separation is less than a fixed minimum peak distance (*d*_peak_ = 3 ms, i.e. fewer than *d*_peak_ × *f*_*s*_ samples apart) are merged and counted as a single crossing. Crossings separated by more than *d*_peak_ are counted independently. The peak distance is not exposed in the graphical interface but can be modified in the source code to suit different target taxa. Triggering occurs when the cumulative number of threshold crossings within the preceding pre-trigger window equals or exceeds a user-defined minimum (default 2). Upon triggering, a tap-mode clip is captured and a configurable cooldown period (default 0.25 s) is imposed to suppress immediate re-triggering. Effective use of the auto-trigger requires some knowledge of the target sound pattern: the minimum crossing count should be set high enough to reject transient noise yet low enough to capture the vocalisations of interest.

If a calibration factor *C*_Pa_ (Pa per input unit) is provided, the threshold in decibels is converted to an amplitude threshold in device units via 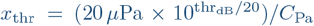 In the absence of calibration, the threshold is interpreted as dBFS and converted directly via 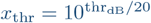.

Figure 4 summarises the ring-buffer layout and the auto-trigger decision logic. The fixed peak distance of 3 ms is chosen to resolve individual pulses in bat echolocation sequences; for other taxa, a crossing count of 1 or 2 with an appropriate amplitude threshold is usually sufficient.

### 2.6 Calibration and sound pressure estimation

The calibration constant *C*_Pa_ (Pa per input unit) is user-adjustable at all times and does not affect the acquisition or recording pipeline. When specified (*C*_Pa_ *>* 0), waveform display levels are converted to sound pressure level (SPL) according to:

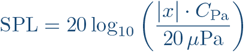

with a floor of 0 dB SPL and the waveform axis labelled in “dB re 20 µPa” over the range [0, 140] dB. In the absence of calibration, amplitude is displayed in dBFS relative to full scale:

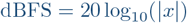

with a floor of −120 dBFS and the axis spanning [−120, 0] dB. Calibration state transitions are tracked internally so that axis labels and limits are updated only when the calibration mode changes.

The calibration factor also governs the interpretation of auto-mode thresholds (Section 2.5), enabling the users to configure triggering in absolute SPL when a calibrated microphone is available, or in relative dBFS during uncalibrated exploratory surveys.

### 2.7 Monitoring and heterodyne processing

Real-time monitoring supports three modes: disabled (*Off*), passthrough, and heterodyne. The monitoring path operates independently of the recording path and does not modify stored data. In heterodyne mode, the input signal is frequency-shifted by multiplication with a cosine carrier at a user-defined frequency (1–(*f s/*2) kHz) with phase-continuous accumulation across frame boundaries—the same technique described in detail for ESPERDYNE (Umadi, 2026a) and BATSY4-PRO (Umadi, 2025b). Output channels support one-to-one mapping, explicit left-/right ear assignment, or an odd/even mix mode. A software gain (0–4×) is applied, followed by hard clipping to ±1.

When input and output sampling rates differ, rational resampling (`resample()` if the Signal Processing Toolbox is available, otherwise integer decimation) is applied per column. Output samples are buffered and drained in fixed-size chunks matching the writer’s expected frame size, preventing PortAudio frame-size mismatch errors.

### 2.8 Spectral analysis and visualisation

Two independent live spectrograms (channels A and B) are computed via short-time Fourier transform using a periodic Hann window of 1 024 samples, zero-padded to an FFT length of 2 048, with no overlap (hop size equal to the window length). Power spectral density columns are expressed in decibels (20 log_10_(|*X*| + 10^−9^)) and stored in single-precision circular buffers. The number of display columns is determined by

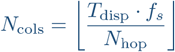

where *T*_disp_ is the display duration (set to 3.0 s). Each spectrogram buffer has an independent write index, and the circular contents are linearised before being pushed to the display.

The spectrogram buffers are updated at every audio frame (i.e. at the full audio frame rate), whereas graphical rendering occurs only at the user-configurable UI refresh rate (default 15 Hz, adjustable from 1 to 60 Hz). This decoupling ensures that pixel-level drawing operations do not introduce latency into the acquisition loop. The effective maximum useful frame rate is bounded by ⌊*f*_*s*_/*N*_frame_⌋.

The spectrogram upper frequency limit defaults to *f*_*s*_/2 and may be dynamically restricted to focus on biologically relevant frequency bands (e.g. bird calls). A greyscale colourmap is used with a dynamic range of [−120, 0] dB.

### 2.9 WAV file writing

Audio files are written in standard RIFF/WAVE format. Tap-mode recordings (Section 2.5) use MATLAB’s built-in `audiowrite()` function. Continuous and auto-mode recordings employ a custom low-level streaming writer that operates via three stages:

1. *Open*: A standard 44-byte RIFF/WAVE header is written with format code 1 (PCM) or 3 (IEEE 754 float). The RIFF and data chunk size fields are initialised as zero placeholders.
2. *Write*: Incoming samples are clipped to [−1, 1] and written as raw bytes at the selected bit depth: `int16` scaling for 16-bit, three-byte little-endian packing for 24-bit, `int32` scaling for 32-bit integer, or `single` for 32-bit float.
3. *Close*: The file pointer seeks back to byte offsets 4 and 40 to patch the RIFF and data chunk sizes, ensuring compliance with standard WAV readers.

The bit depth is selectable by the user and is honoured across all recording modes. Output filenames follow the convention `yyyyMMdd_HHmmss_SSS[_suffix].wav,` and files are written to a default directory structure`(∼/Documents/RUBAT/Rec/YYYYMMDD/)` that may be overridden by the user.

### 2.10 Portability and compilation

RUBAT Studio v4.0 was packaged using the MATLAB Application Compiler to produce standalone executables requiring only the freely distributable MATLAB Runtime. The software has been tested on macOS (Core Audio) and Windows (WASAPI/ASIO via PortAudio abstraction). Platform-specific adaptations include asynchronous port release handling on macOS and font resolution across operating systems.

The Audio Toolbox is the sole required toolbox dependency; the Signal Processing Toolbox is recommended for high-quality rational resampling but is not essential, as integer decimation is used automatically when it is unavailable. No proprietary hardware or vendor-specific drivers are required beyond those needed for the selected audio interface, making RUBAT Studio compatible with any standard external or built-in sound device.

### 2.11 Testing and field recordings

RUBAT Studio was tested with a range of audio interfaces: RME Babyface Pro (up to 192 kHz), Focusrite Scarlett 2i2, Behringer UMC404HD, Dodotronic Ultramic 384K (384 kHz), and the built-in sound card of a MacBook Pro. Through Apple Continuity, the software also recognised an iPhone 13 Pro and AirPods Pro as audio devices. Multi-hour continuous runs on both macOS and Windows completed without crashes or artefacts. Figure 5 shows spectrograms from a two-channel field recording made on the Weihenstephaner Berg in Freising, Germany, in early spring 2026 using a RODE NT5 and a Sanken CO-100K microphone via an RME Babyface Pro at 192 kHz.

**Figure 5.**
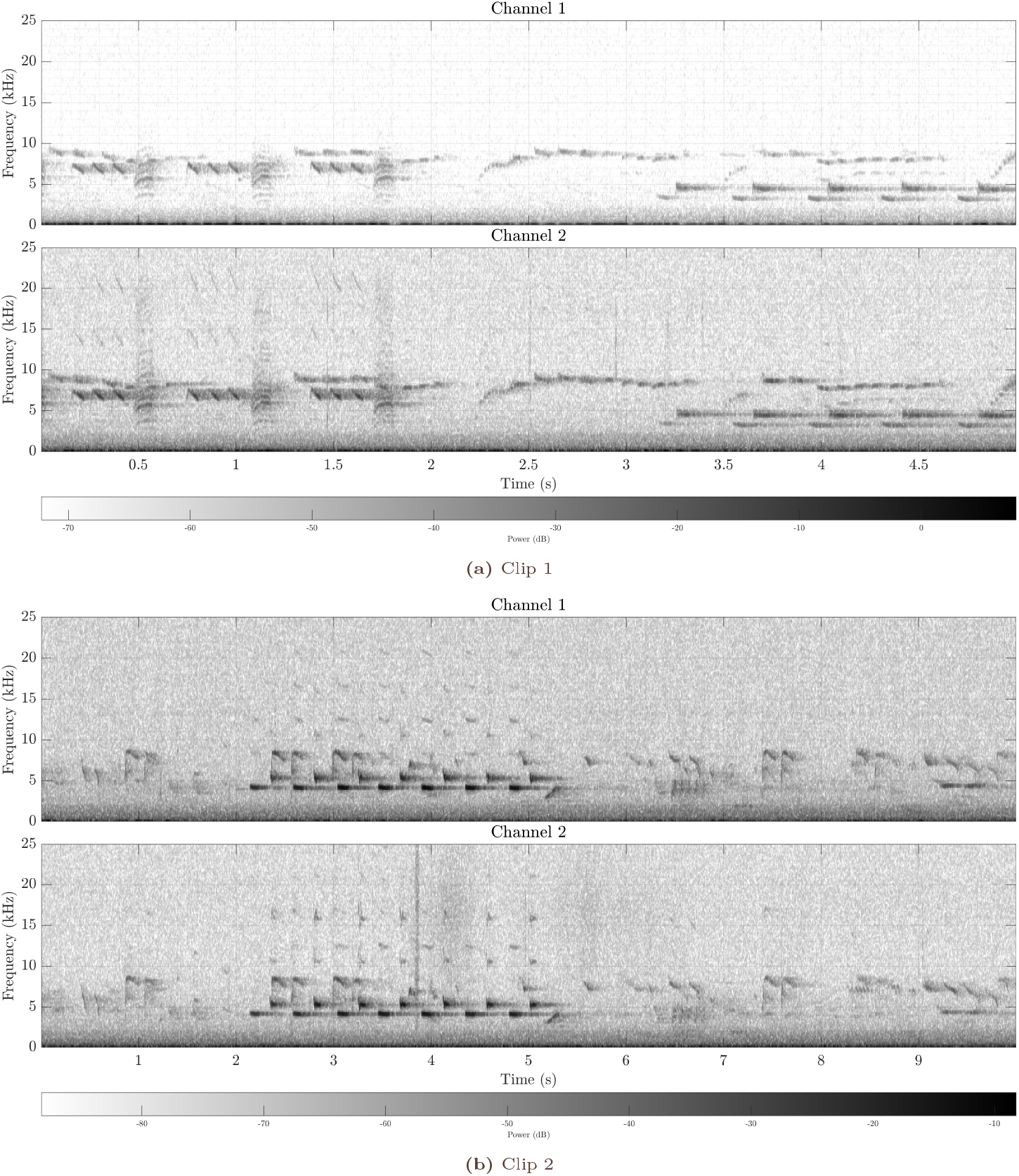
Two-channel spectrograms from field recordings made on the Weihenstephaner Berg, Freising, Germany, in early spring 2026. Channel 1: RODE NT5; Channel 2: Sanken CO-100K. Audio was captured at 192 kHz via an RME Babyface Pro. Power is displayed on a greyscale (darker = higher power).

## 3 DISCUSSION

RUBAT Studio addresses a persistent gap in the bioacoustic data-collection workflow: the absence of an accessible, hardware-agnostic, multichannel recording application that integrates the device management, real-time monitoring, calibration, and automated acquisition features required for full-band field and laboratory research. By consolidating these capabilities into a single interface, RUBAT lowers the technical barrier to multichannel recording and makes the workflow accessible to operators without requiring high-level audio engineering or platform-specific system configuration expertise.

### 3.1 Unified device control and technical accessibility

By exposing the full set of audio device parameters — sampling rate, bit depth, buffer size, and channel selection — in a single panel, RUBAT eliminates the need to navigate operating system preferences, proprietary driver utilities, or, most importantly, *ad hoc* scripts. Once hardware is connected, the interface guides the user through device probing, channel assignment, monitoring configuration, and recording—a workflow that previously required disparate tools and platform-specific knowledge. Students and early-career researchers can therefore be trained rapidly. Real-time waveform and spectrogram feedback, with selectable heterodyne or passthrough monitoring, further assists in setup verification before a session begins.

### 3.2 Laboratory multi-setup operation

In laboratory settings, RUBAT can serve as a central recording terminal for multiple concurrent experimental configurations. For instance, consider a facility operating four acoustic chambers, each equipped with a dedicated multichannel audio interface for recording animal vocalisations (Figure 2). All four interfaces can be connected to a single host computer running RUBAT; the experimenter selects the desired device, channels, and recording parameters from the interface without physically disconnecting or reconnecting hardware. This single-terminal paradigm simplifies logistics, reduces setup time between experiments, and eliminates a common source of configuration errors.

### 3.3 Calibration and signal-level assurance

The real-time calibration feature warrants particular emphasis. Sound-pressure-level calibration of recording systems is a technically demanding multi-step procedure—typically involving a reference tone source, a calibrated reference microphone, and post-hoc gain correction. RUBAT simplifies this process to a single parameter: the calibration constant *C*_Pa_ (Pa per input unit; see Section 2.6). When set, the waveform display presents levels in dB SPL (re 20 µPa) and the automatic trigger (Section 2.5) interprets thresholds in absolute SPL, enabling ecologically meaningful detection criteria. Validation is equally straightforward: a known reference tone can be played at any time, and the displayed level checked against the expected value to confirm that the calibration holds. This built-in validation pathway supports reproducible and traceable data collection, which is essential when quantitative sound-pressure measurements are required—for example, in source-level estimation from calibrated microphone arrays (de Framond et al., 2023; Madsen & Wahlberg, 2007).

### 3.4 Open source and cost considerations

The open-source distribution model of RUBAT Studio (v4.0) carries several advantages. First, the full source code is available upon request for inspection, modification, and extension, enabling research groups to adapt the software to specialised workflows without vendor dependence. Second, RUBAT Studio (v4.0) bears no licensing cost. Commercial ultrasound recording solutions such as Avisoft RECORDER or BATSOUND carry licence fees that, when combined with the cost of the required proprietary hardware, can place multichannel acoustic setups beyond the budget of many research groups—particularly those in low- and middle-income countries where bat diversity and conservation needs are often greatest (Manzano-Rubio et al., 2022). RUBAT, together with open-source hardware platforms such as BATSY4-PRO (Umadi, 2025b) and ESPERDYNE (Umadi, 2026a), offers a fully open alternative that aligns with the broader movement toward open-source scientific instrumentation (Pearce, 2015).

### 3.5 Integration with downstream analysis pipelines

RUBAT records standard RIFF/WAVE files at user-selected bit depths (16, 24, or 32-bit PCM and 32-bit float; Section 2.9), ensuring compatibility with virtually all bioacoustic analysis software. Recorded data can be channelled directly into sorting and quality-control tools such as BatReviewer (Umadi, 2025a), spectral analysis environments such as Sonic Visualiser and Raven Pro, or automated deep-learning classifiers such as BatDetect2 (Aodha et al., 2022). When paired with array geometry optimisation tools such as WAH-*i* (Umadi, 2026b; Umadi, 2025c), RUBAT supports a complete pipeline from optimised sensor placement through multichannel acquisition to automated species identification and acoustic localisation.

### 3.6 Real-time output and cross-species behavioural experiments

Because RUBAT routes acquired audio to a selectable output device in real time (Section 2.7), vocalisations captured on one interface can be delivered through loudspeakers or ultrasonic transducers on a second interface, enabling experiments in which animals are acoustically coupled but visually isolated. For example, natural echolocation calls from a freely flying bat can be streamed to moths in a separate observation chamber, preserving the full behavioural variability of a hunting bat—call-by-call adjustments in level, bandwidth, and pulse interval—that synthetic playback cannot reproduce. The same paradigm extends to songbird vocal learning or any scenario requiring acoustic stimulus delivery with visual-cue exclusion, offering a flexible framework for dissecting sensory communication across taxa.

### 3.7 Broader applicability and outlook

Although RUBAT was motivated by bat bioacoustics, the software imposes no taxon-specific assumptions. Any application requiring multichannel high-sample-rate recording with real-time monitoring—bioacoustic surveys of birds, insects, or marine mammals (via hydrophone arrays), industrial ultrasound inspection, or acoustic-scene capture for soundscape ecology (Pijanowski et al., 2011; Schoeman et al., 2022)—can use RUBAT with appropriate hardware.

### 3.8 Extensibility and future directions

The software architecture enables several extensions that would leverage the existing device-management and streaming infrastructure. Real-time machine-learning classifiers could be embedded in the capture loop for on-the-fly species identification and automated dataset construction. The multichannel acquisition core could also be coupled with beamforming and source-localisation routines to track sound sources and reconstruct three-dimensional flight trajectories from microphone-array recordings. Each extension requires adding per-frame signal-processing routines to the existing capture loop.

In summary, RUBAT Studio fills a critical niche in the bioacoustic recording software landscape. By providing a unified, free, cross-platform interface that combines multichannel device management, real-time acoustic and visual monitoring, automated threshold-triggered recording, and single-parameter calibration, it significantly lowers the barrier to rigorous bioacoustic data collection—whether in a controlled laboratory with multiple experimental setups or in the field with portable multichannel hardware. As microphone-array methods and automated analysis pipelines continue to advance, the availability of robust, accessible, and extensible recording software becomes increasingly important for translating technological progress into ecological and conservation outcomes.

## ACKNOWLEDGEMENTS

I thank Uwe Firzlaff and colleagues at the TUM for their support and institutional assistance throughout the development of this work.

## COMPETING INTERESTS

The author declares no competing interests.

## ETHICAL STATEMENT

This work describes the design and implementation of a software tool for bioacoustic recording. No animal handling, experimentation, or data collection from live animals was involved; therefore, no ethical approval was required.

## DATA & CODE AVAILABILITY

The RUBAT Studio source code and standalone installers are available on GitHub at https://github.com/raviumadi/RUBAT-Studio.

User guide and documentation are published at https://raviumadi.github.io/RUBAT-Studio.

## FUNDING

This work was not externally funded.

## Notes

https://rubat.biosonix.io

### Competing Interest Statement

The authors have declared no competing interest.

